# The gut microbiota regulates mouse biliary regenerative responses

**DOI:** 10.1101/575050

**Authors:** Wenli Liu, Chao Yan, Bo Zhang, Renjin Chen, Qian Yu, Xiangyang Li, Yuzhao Zhang, Hui Hua, Yanxia Wei, Yanbo Kou, Zhuanzhuan Liu, Renxian Tang, Kuiyang Zheng, Yugang Wang

## Abstract

Injury to the biliary epithelium triggers cholangiopathies. However, factors involved in regulating the pathogenesis remains incompletely understood. Here, we report that the gut microbiota is important in regulating hepatobiliary fibroproliferative regenerative process. In helminth *Clonorchis sinensis*-induced bile duct injury model, wild-type mice showed more extensive peribiliary fibroproliferative responses than non-littermate IL-33-deficient mice. However, these reactions could be attenuated by co-housing of the animals together. In the meantime, the relatively fibroproliferative-resistant IL-33-deficient mice could become fibroproliferative-responsive especially in large intrahepatic bile duct by antibiotics treatment. Furthermore, microbiota-derived metabolite butyrate was able to inhibit biliary organoid expansion *in vitro* and temper 3,5-diethoxycarbonyl-1,4-dihydrocollidine-induced biliary fibrosis *in vivo*. Together, our data implies a potential way of management of hepatobiliary diseases by modulating gut microbiota.

## INTRODUCTION

The biliary tree including intrahepatic and extrahepatic bile ducts (IHBD and EHBD) forms the primary path by which bile is secreted by the liver then transported to the gut. Disruption of the epithelial lining of bile ducts triggers hepatobiliary inflammation, fibrosis, and even malignant transformation, which collectively were referred as cholangiopathies. The cholangiopathies account for substantial morbidity and mortality and are often progressive that lead to end-stage liver disease. Despite the severe disease phenotypes, the molecular and cellular effectors and their relationships to cholangiopathies remain poorly defined ^1^.

A number of animal models have been generated to provide in vivo tools to investigate the mechanisms of biliary repair and, eventually, to test the efficacy of innovative treatments ^2^. Each of the animal model has its own limitations in translating to human settings. 3,5-Diethoxycarbonyl-1,4-dihydrocollidine (DDC)-feeding in mice recapitulated pathogenic hallmarks of human primary sclerosing cholangitis (PSC) ^3^. *Clonorchis sinensis* is a food-borne helminth that naturally cause cholangiopathies ^4, 5^. During progression of biliary injury induced by the worm, a disorganized expansion of bile ducts and recruitment of inflammatory cells known as ductular reaction (DR) become so prominent that not only biliary epithelial but also the peribiliary glands (PBGs), clusters of epithelial cells residing in the submucosal compartment of the bile duct, show hyperplasia and proliferation. PBGs participate in the secretion of neutral, carboxylated, and sulphated mucins into the bile duct lumen. The term adenomatous hyperplasia or chronic proliferative cholangitis has been used to describe this phenomenon ^6^. What is the underlying molecular and cellular mechanisms of these biliary responses remains incompletely understood.

The gut microbiota can influence or modulate chronic liver disease ^7^. In the absence of the intestinal microbiota, germ-free *mdr2^-/-^* mice had exacerbated primary sclerosing cholangitis ^8^, while germ-free NOD.c3c4 mice had attenuated spontaneous biliary inflammation ^9^. Whether the gut microbiota participates in the hepatobiliary pathogenesis during *C. sinensis* infection is largely unknown.

Our recent study showed significantly increased biliary epithelial IL-33 expression and serum IL-33 levels in patients with *C. sinensis* infection ^10^. Therefore, we were initially interested in a potential role of IL-33 in the cholangiopathies during *C. sinensis* infection. Unexpectedly, we found that the gut microbiota instead played important roles in regulating helminth injury-induced hepatobiliary regenerative responses. We further showed that microbiota-derived metabolite butyrate was able to inhibiting biliary tree stem/progenitor cell expansion in vitro and tempering biliary fibrosis during 3,5-diethoxycarbonyl-1,4-dihydrocollidine intoxication in vivo. Thus, gut microbiota is an important regulator of hepatobiliary fibroproliferative regenerative responses.

## RESULTS

### WT mice have more extensive hepatobiliary fibroproliferative responses compared to non-littermate IL-33-deicient mice

Our initial goal is to study the role of IL-33 during helminth infection, so we generated germline *IL-33^-/-^* mice (Fig. S1). We intragastically infected non-littermate 6- to 8-week-old wild-type (WT) and *IL-33^-/-^* mice with *C. sinensis* metacercariae to induce biliary inflammation, and then monitored the hepatobiliary fibro-inflammatory changes at day 35 post infection. Based on histopathological evaluation of H&E-stained sections, we found extensive ductular reaction in the non-littermate WT mice both inside the liver (Fig. 1A) and in the EHBD (Fig. 1B), which were all attenuated in *IL-33^-/-^*. We next examined liver fibrosis by evaluation of Masson Trichrome-stained sections and observed a significantly higher percentage of fibrotic area in the non-littermate WT mice, which was also alleviated in the *IL-33^-/-^* mice (Fig. 1C). We also assessed fibrosis by biochemical measurement of hepatic hydroxyproline (hyp) concentration (Fig. 1D). The hepatic hyp levels were initially similar between uninfected naïve WT and *IL-33^-/-^* mice, 35 days post infection the hyp concentration increased more than 3 folds in the WT and less than 1-fold in the *IL-33^-/-^* mice. Among the infected mice, there were about 2-3 folds reduction of the hyp levels in the *IL-33^-/-^* compared to WT mice, indicating fewer collagen deposition and therefore less severe fibrotic response in the *IL-33^-/-^* mice.

**Figure 1.**
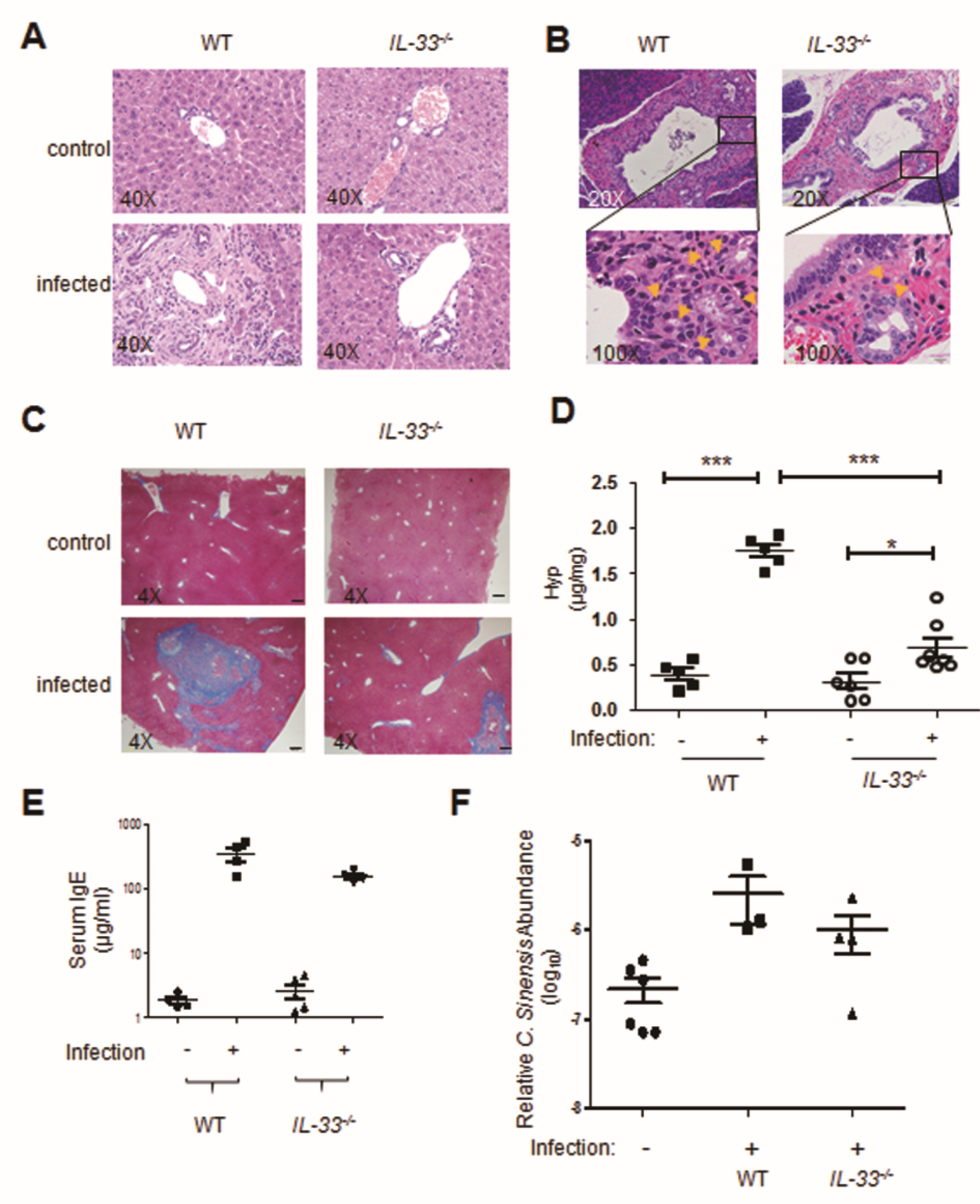
WT mice have more extensive hepatobiliary fibrosis compared to non-littermate IL-33-deicient mice. **(A)** Representative images of H&E-stained liver sections of uninfected control and *C. sinensis*-infected WT and *IL-33^-/-^* mice (n=5 or 6). (scale bar, 20 µm.) **(B)** Representative H&E-stained images of the EHBD of infected WT and *IL-33^-/-^* mice. Arrows indicate peribiliary glands. Upper panel, scale bar = 50µm; lower panel, scale bar = 10 µm. **(C)** Representative images of Masson’s Trichrome-stained liver sections of uninfected and *C. sinensis*-infected WT and *IL-33^-/-^* mice. (Scale bar, 200 µm.) **(D)** Concentrations of hydroxyproline in the hepatic homogenate of control and *C. sinensis*-infected mice. Each symbol represents one mouse. **(E)** Total serum IgE production from the indicated mice. **(F)** Relative abundance of *C. sinensis* in the liver tissues of the indicated mice at 35 days post-infection quantified by qPCR. \*\**P* < 0.01, \*\*\**P* < 0.001.

In contrast to the attenuated fibrotic response, the serum IgE responses were not attenuated in the *IL-33^-/-^* compared with WT mice (Fig. 1E). This is consistent with the notion that IL-33 signaling is not absolutely required for IgE responses during helminth infection ^11^. The parasite burden was also comparable between these two strains of mice (Fig. 1F), ruling out potential antigenic load difference in dominating the different fibrotic responses.

### WT mice cohoused with *IL-33^-/-^* mice have reduced fibroproliferative responses comparing to WT mice cohoused with WT mice

One caveat of our experiment is that we used non-littermate WT and *IL-33^-/-^* mice to compare their fibrotic difference during helminth infection. To rule out a potential role of microbiota in influencing the outcome, we co-housed the non-littermate WT with *IL-33^-/-^* mice (refer to as WT(*IL-33^-/-^*) and *IL-33^-/-^*(WT), respectively) for 5 weeks starting at day 0 after helminth infection. Based upon unweighted UniFrac principal coordinate (PCoA) analysis of the cecal bacterial 16S rRNA genes, the structures of the bacterial communities in the cohoused WT(*IL-33^-/-^*) and their cagemates *IL-33^-/-^*(WT) mice became more similar than those in the WT(WT) and *IL-33^-/-^*(*IL-33^-/-^*) mice (Fig 2A). To our surprise, in comparison with WT mice that co-housed with WT mice (refer to as WT(WT)) mice, disease severity reduced significantly in the WT(*IL-33^-/-^*) mice (Fig. 2B and 2C). The liver fibrosis in the WT(*IL-33^-/-^*) became very mild and was comparable to that of their cagemates *IL-33^-/-^*(WT) mice (Fig. 2B and 2C). The results indicate a potential previously unappreciated role of the gut microbiota in regulating peribiliary fibroproliferative responses during *C. sinenesis* infection.

**Figure 2.**
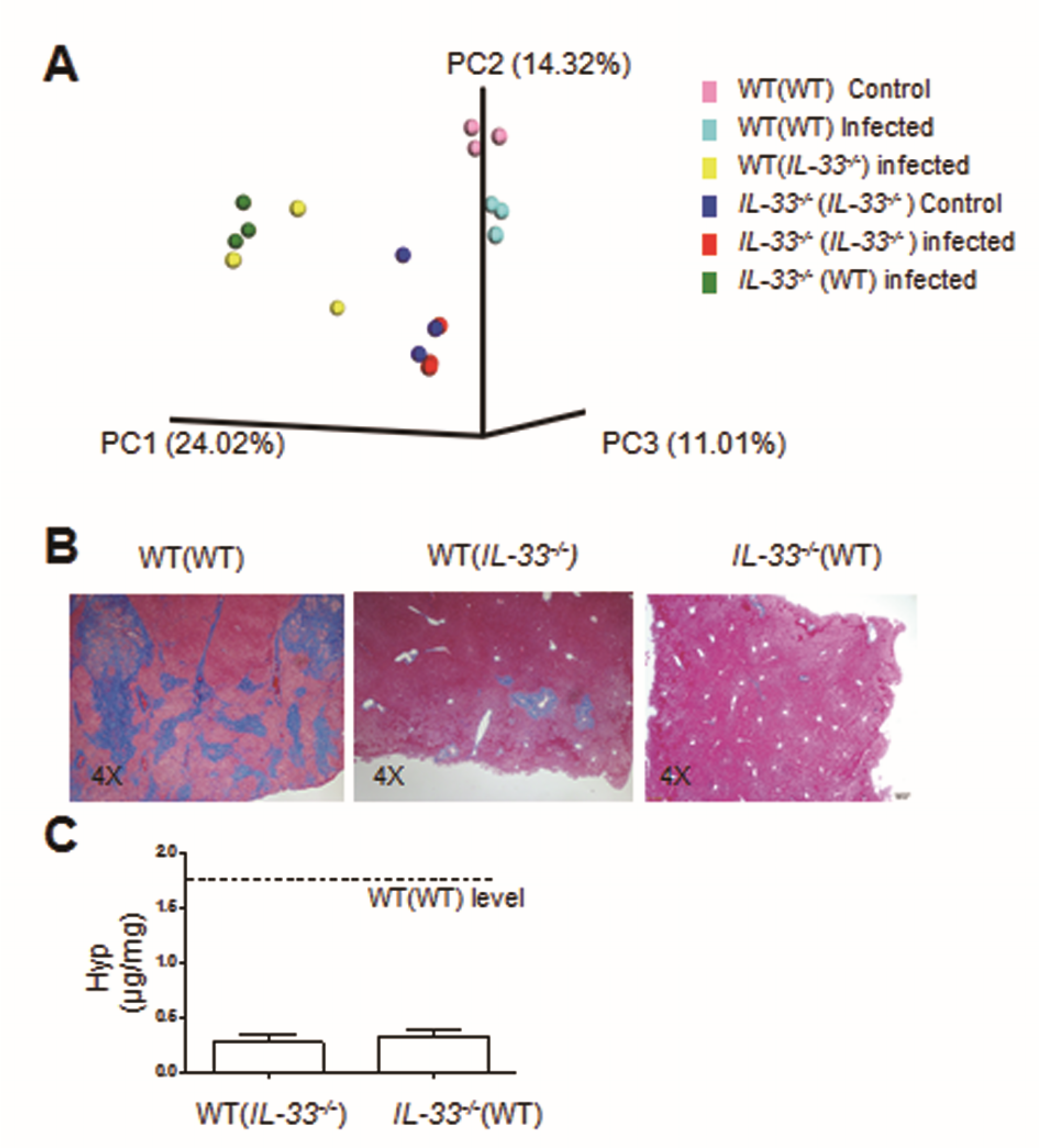
WT mice cohoused with *IL-33^-/-^* mice have reduced fibroproliferative responses comparing to WT mice cohoused with WT mice. **(A)** Unweighted UniFrac principal coordinate analysis according to cecal bacterial 16S rRNA sequences showing microbiota compositional differences among naïve WT(WT) and *IL-33^-/-^* (*IL-33^-/-^*), infected WT(WT) and *IL-33^-/-^* (*IL-33^-/-^*), and infected *WT(IL-33^-/-^) and IL-33^-/-^ (WT)* mice. Each symbol represents one mouse. **(B)** Representative images of H&E-stained liver sections of the indicated mice that infected by *C. sinensis* (n=5 or 6). **(C)** Hepatic hydroxyproline concentrations in the cohoused WT and IL-33-/-mice (n=5 or 6).

### Microbiota involves in hepatobiliary fibroproliferative regenerative response during helminth infection

To test for a potential role of microbiota in regulating biliary fibroproliferative response during helminth infection, we fed both WT and *IL-33^-/-^* mice with a broad-spectrum cocktail of antibiotics (Abx) in the drinking water *ad libitum* during *C. sinensis* infection. Unweighted UniFrac PCoA analysis indicated that the microbiota compositions changed dramatically after Abx treatment (Fig. S2A). The bacterial diversity was greatly diminished after Abx treatment, the firmicutes and bacteroidetes were mostly gone and the remaining left bacteria were dominantly proteobactria (Fig. S2B).

We next investigated whether there were differences in biliary fibrosis after antibiotics treatment. Based on histopathological evaluation of Masson Trichrome-stained sections, more severe peribiliary fibrotic response was found in the antibiotics-treated *IL-33^-/-^* mice compared to the antibiotics-untreated *IL-33^-/-^* mice (Fig. 3A and 3B). The peribiliary fibroproliferative changes were largely confined to large IHBD (Fig 3A). The overall hepatic hyp levels were not increased after Abx treatment (data not shown). To test whether antibiotics treatment might influence the helminth egg production, we examined the whole liver *C. sinensis* egg number microscopically by precipitation the whole liver eggs through 70% percoll density gradient. There was no obvious egg number difference between untreated and antibiotics-treated *IL-33^-/-^* mice (30±26 vs.11±11). The data collectively suggest that microbiota involves in hepatobiliary fibrotic regenerative responses.

**Figure 3.**
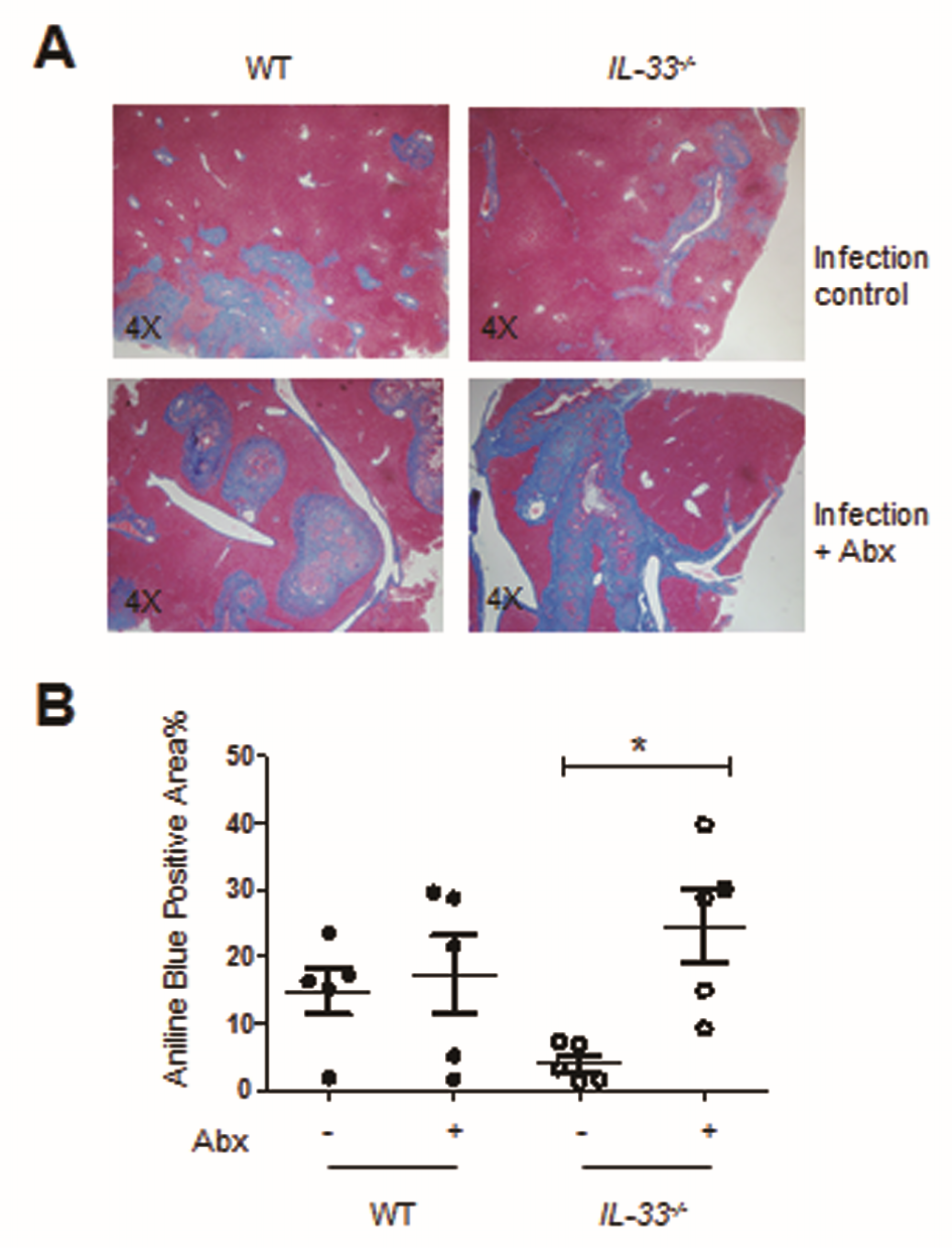
Gut microbiota regulates hepatobiliary fibrotic regenerative response. **(A and B)** mice were treated with a broad-spectrum antibiotics cocktail during helminth infection. (A) Representative images of Masson’s Trichrome-stained liver sections of the antibiotics-treated and untreated WT and *IL-33^-/-^* mice (n=5 or 6). Scale bar, 200 µm. (B) Quantitation of collagen deposition by densometric scanning. Each symbol represents one mouse. \**P* < 0.05.

### Abx treatment unleashes strong hepatobiliary regenerative response after tissue injury in IL-33-deficient mice

To determine the potential mechanisms by which the microbiota influences peribiliary fibrosis during clonorchiasis, we performed transcriptional profiling analysis of liver RNAs isolated from antibiotics treated and untreated WT and *IL-33^-/-^* mice 35 days post helminth infection. There were 292 hepatic genes significantly upregulated and 477 hepatic genes downregulated more than 2-fold after Abx treatment in *IL-33^-/-^* mice (Fig. 4A). KEGG pathway analysis of the differential-expressed genes between antibiotics treated and untreated *IL-33^-/-^* mice unveiled that Abx treatment in *IL-33^-/-^* mice perturbed host-microbiota interaction pathway (e.g. Toll-like receptor signaling pathway), influenced hepatic metabolic functions (e.g., steroid biosynthesis, xenobiotics metabolism, bile secretion etc.), and induced extracellular matrix gene expressions (e.g. *Tnxb, VWF, Col6a1, Col6a2, Col6a5* and *Lamb2*) that are involved in fibrotic responses (Fig. 4B). We also found a few common genes that were upregulated more than 2-fold in both antibiotics-treated WT and *IL-33^-/-^* mice (Fig. 4C). These genes, including *sox9, Ihh, muc6, tff2, mmp7* and *atg*, have all been implied to be involved in either ductular regeneration or cholangiocyte proliferation and activation ^12–19^, suggesting potential molecular pathways of microbiota in regulating biliary regenerative programs.

**Figure 4.**
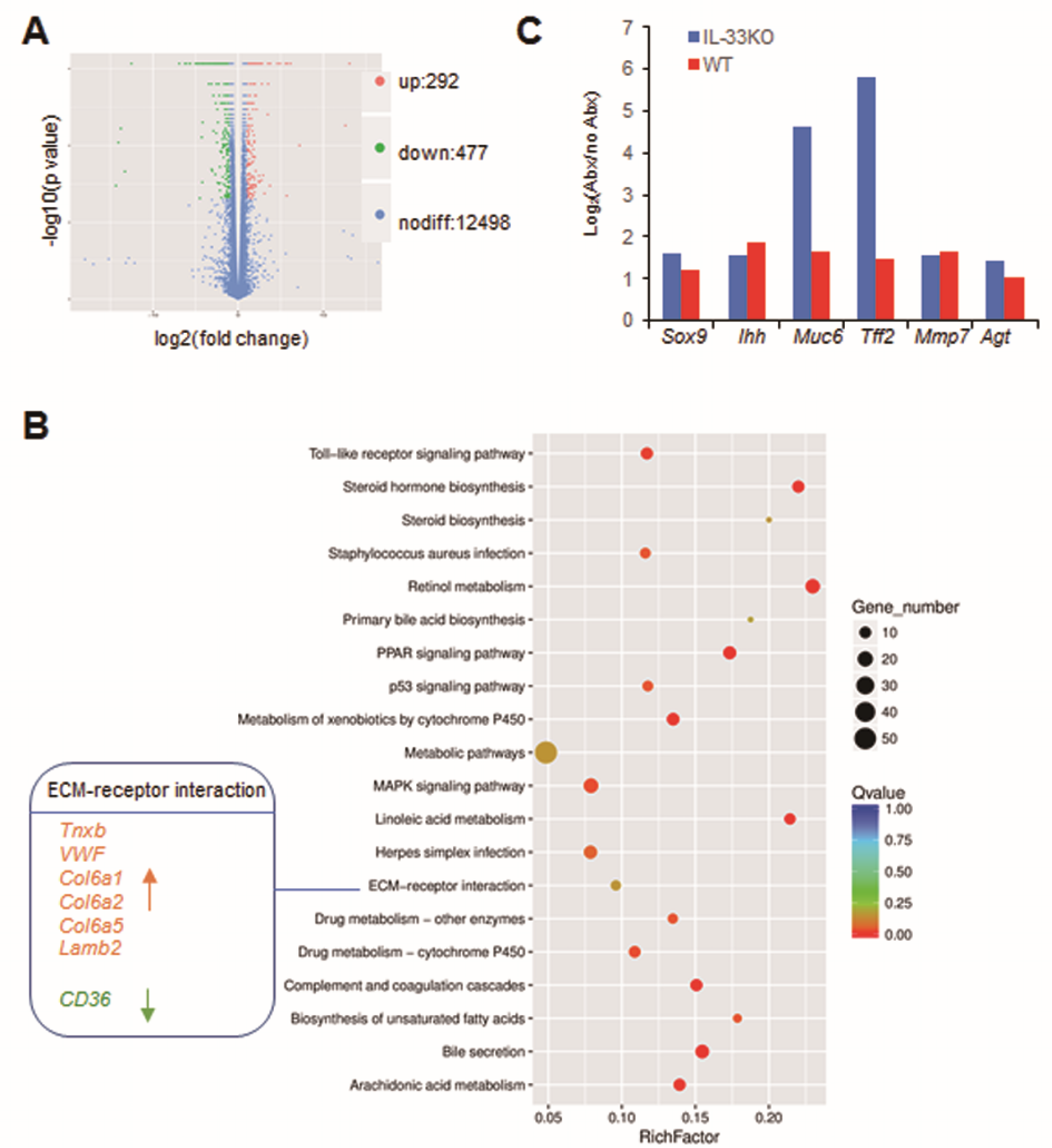
Abx treatment unleashes strong hepatobiliary regenerative response after tissue injury in IL-33-deficient mice. **(A)** Volcano plot showing hepatic differential expressed genes before and after antibiotics treatment in *C. sinensis*-infected IL-33-deficient mice. **(B)** Top 20 statistical significantly enriched liver KEGG pathways for antibiotics-treated vs. untreated *C. sinensis*-infected IL-33-deficient mice. **(C)** List of the shared genes between antibiotics-treated WT and *IL-33^-/-^* mice, all of which were statistical significantly upregulated more than 2-fold.

### Gut microbiota-derived metabolite butyrate inhibits biliary organoid expansion

Since increased *sox9* gene expression after antibiotics treatment suggests of cholangiocyte expansions, which are in tightly controlled by biliary stem cell (BTSC) activity. We suspected that microbiota might be able to regulate BTSC. A prominent bacterial metabolite butyrate has been proved to be a potent inhibitor of intestinal stem/progenitor cell proliferation ^20^, we thus posited that butyrate might also be able to regulate BTSC during tissue injury since BTSC and intestinal stem cells are developmentally derived from the same source ^15, 21^. Abx treatment can influence butyrate production. It not only wiped out main butyrate-producing bacteria estimated by the butyryl-CoA CoA transferase gene (Fig. 5A), the butyrate concentration in the intestinal lumen was also reduced significantly (Fig. 5B). To prove a role of butyrate in regulating BTSC function, we established biliary organoid culture. Consistent with a role of IL-33 in mediating biliary epithelial proliferation ^22^, rmIL-33 promoted biliary organoid expansion indicated by Ki67 immunostaining (Fig. 5C and 5D). More importantly, butyrate was able to inhibit BTSC expansion both in the presence or absence of rmIL-33 (Fig. 5C and 5D). Thus, microbiota-derived butyrate has the potential to participate in regulating biliary regenerative program at least in part via inhibiting BTSC bioactivity.

**Figure 5.**
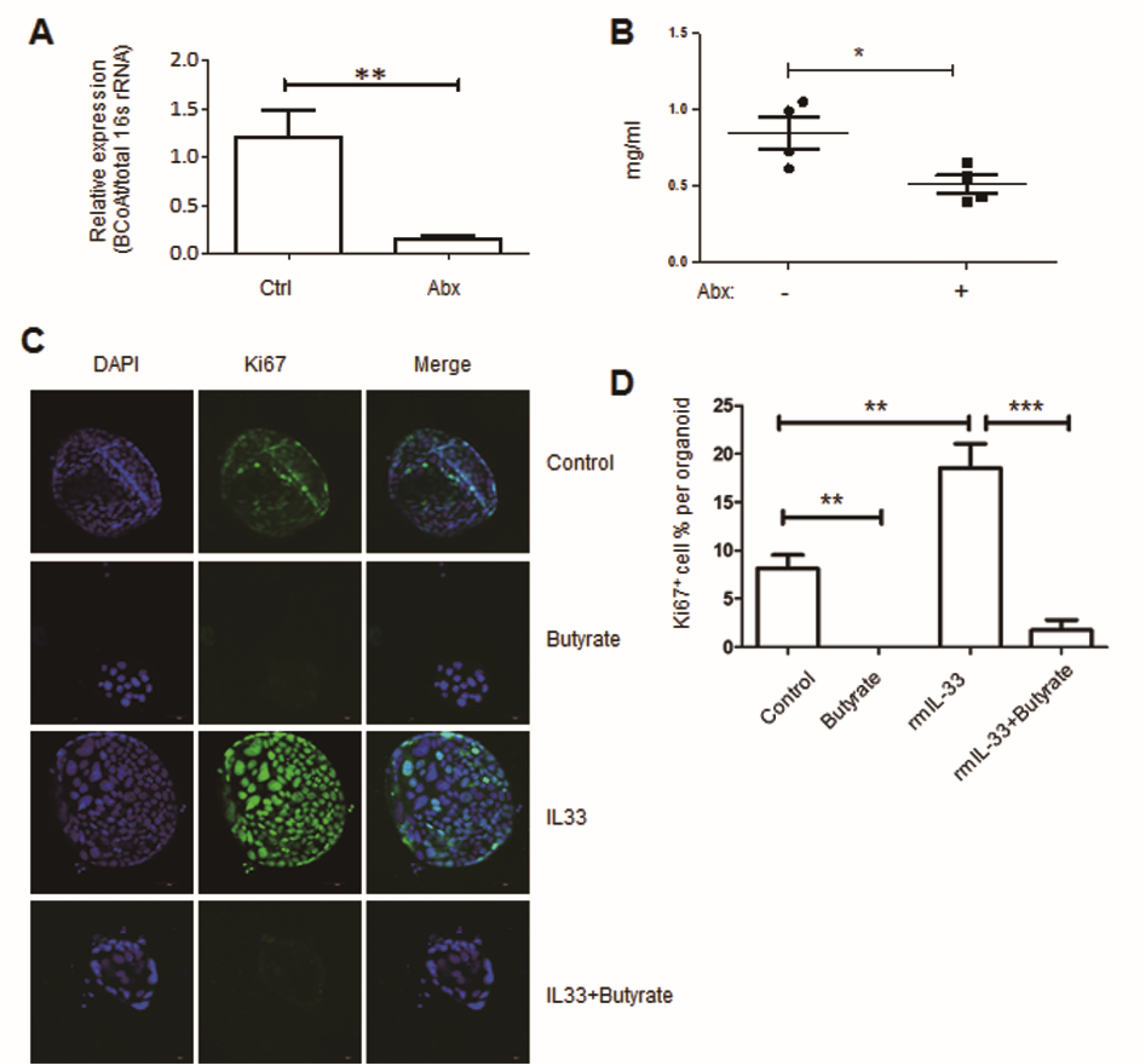
Butyrate inhibits biliary organoid expansion. **(A)** Relative expression of butyryl-CoA CoA transferase gene in cecal contents after abx treatment. **(B)** Butyrate concentration in cecal contents determined by LC-MS. **(C)** Representative confocal microscopic images of biliary organoids that treated with indicated reagents (magnification, 20X). **(D)** Quantitation of the percentages of Ki67 positive cell number among DAPI-stained positive cells. \*\**P* < 0.01, \*\*\**P* < 0.001.

### Butyrate inhibits biliary fibrosis *in vivo*

In order to test whether butyrate has a role in inhibiting biliary fibrosis *in vivo*, we switched to a well-established 3,5-diethoxycarbonyl-1,4-dihydrocollidine (DDC)-induced sclerosing cholangitis and biliary fibrosis model ^3^. WT or *IL-33^-/-^* mice were fed a 0.1% DDC-supplemented diet and with or without 10 g/L sodium butyrate in the drinking water for 4 weeks. Although butyrate did not have any role in protecting hepatobiliary system from DDC-induced injury according to serum liver enzymes and metabolites measurements (Fig. S3A-S3D). However, butyrate was effective in attenuating fibrosis in the WT mice, evaluated by biochemical measurement of hepatic hydroxyproline concentration (Fig. 6). The data together indicated that gut microbiota metabolite butyrate can regulate hepatobiliary fibrosis during hepatobiliary injury.

**Figure 6.**
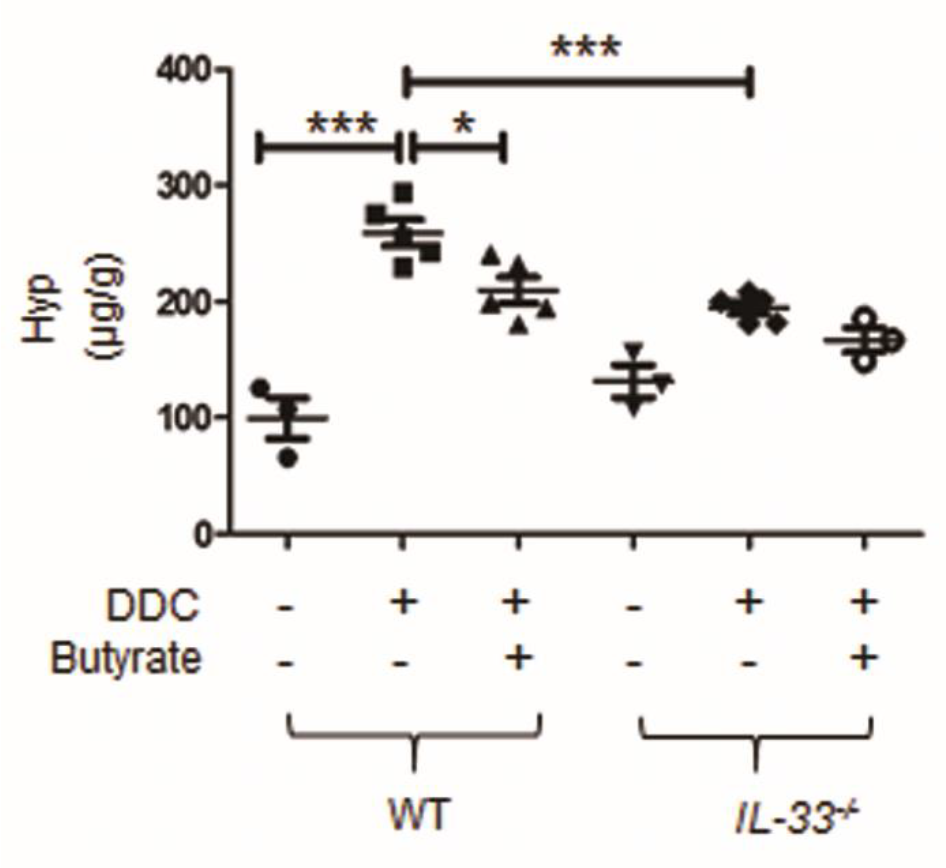
Butyrate inhibits DDC-induced biliary fibrosis. The severity of liver fibrosis induced by 4 weeks of DDC intoxication were evaluated by measurement of the concentrations of hydroxyproline in the hepatic homogenate of the indicated mice. Each symbol represents one mouse. \**P* < 0.05, \*\*\**P* < 0.001.

## DISCUSSION

The cholangiopathies account for substantial morbidity and mortality and are often progressive to end-stage liver disease, however our current understanding of the mechanisms of biliary regenerative process remain limited and innovative treatment has remained to be found. Multiple experimental animal models have been established in the past, most of the methods artificially trigger biliary injury and alter bile acid metabolisms ^2^. In the present study, by using helminth *C. sinensis*- and DDC-induced bile duct injury, we demonstrate that the gut microbiota and their metabolites can play important roles in regulating hepatobiliary fibroproliferative disease progression. Thus, manipulating gut microbiota might hold promise for fibroproliferative hepatobiliary diseases.

Butyrate is a potent inhibitor of intestinal stem/progenitor cell proliferation and wound repair following intestinal injury ^20^. Here we showed that butyrate is also a potent inhibitor of biliary stem/progenitor cells. Butyrate can inhibit BTSC expansion and thus potentially reduce biliary fibroproliferative regenerative responses. However, the cellular and molecular details remain for future investigation. Interestingly recent study suggested a division of labor in hepatobiliary expansion and pathological fibrosis during liver tissue repair ^23^. It is likely that butyrate might involve in regulating multiple tissue repair pathways.

Our current understanding of the impact of microbiota in liver disease is still in its infancy. In a mouse model of primary sclerosing cholangitis, germ-free (GF) *mdr2^−/−^* mice exhibit exacerbated biochemical and histologic features of PSC and increased cholangiocyte senescence, a characteristic and potential mediator of progressive biliary disease ^8^. Aggravation of liver fibrosis was also seen in thioacetamide and carbon tetrachloride (CCl4) treated GF mice compared with thioacetamide- and CCl4-treated conventionally raised mice ^24^. Together with our current data, it seems to be a common theme that the commensals can inhibit epithelium injury-induced fibrotic regeneration programs. While in models of acute hepatic injury such as acetaminophen- and alcohol-induced hepatotoxicity GF mice were protected from hepatic disease ^25, 26^. In a spontaneous biliary inflammation model, GF NOD.c3c4 mice develop a milder biliary affection compared with conventionally raised NOD.c3c4 mice ^9^. It might be that in those models immune-mediated mechanisms were dominant, and the microbiota might be participated in immune cell activation. Regardless which role the microbiota plays in the disease pathogenesis, it is no doubt that gut microbiota is an intriguing regulator in both progression and complications of liver disease.

The role of IL-33 in the cholangiopathies remains to be determined. It has been shown that in a mouse model of extrahepatic cholangiocarcinoma (ECC), biliary epithelial injury-induced regenerative response mediated by IL-33 accelerates development of ECC from peribiliary glands, an effect that was suppressed by anti–IL-33 treatment ^27^. However, the gut microbiota might also play a role in cholangiocarcinoma development, which need to be further explored.

In summary, our data reveals gut microbiota plays important roles in regulating hepatobiliary injury-induced regenerative responses. Manipulating microbiota or its metabolites holds promises in management of cholangiopathies.

## METHODS

### Isolation of *C. sinensis* metacercariae

The metacercariae of *C. sinensis* used in our experiments were isolated from naturally infected *Pseudorasbora parva* collected from clonorchiasis endemic regions in China by digesting the fish with artificial gastric juice (0.7% pepsin in 1% HCl solution, pH2.0). The collections were preserved in cold Alsever’s solution with antibiotics at 4 °C until use.

### IL-33^-/-^ mice

BALB/c wild-type mice were purchased from Shanghai Laboratory Animal Co., Ltd (SLAC, Shanghai, China) and maintained at the SPF facility at Xuzhou Medical University. Germline *IL-33^-/-^* mice were generated by co-microinjection of in vitro translated *Cas9* mRNA and gRNAs into the BALB/c zygotes. The targeting strategy including the gRNA sequences and the knockout allele obtained are depicted in the supplementary information. For parasite infection, 6-8-week-old wild type and *IL-33^-/-^* mice were orally gavaged with 45 metacercariae and monitored daily for diet and survival. For antibiotics treatment, mice were administered ampicillin 1g/l, vancomycin 500 mg/l, neomycin 1g/l and metronidazole 1g/l (AVNM (Sigma-Aldrich China, Shanghai, PR China) in drinking water *ad libitum* the whole time starting from the beginning of the parasite infection. The mice were sacrificed 5 weeks after infection.

### Ethics Statement

All animal experiments were conducted following the guidelines of National Laboratory Animal Ethics Committee of China and performed in accordance with institutional regulations after review and approval by the Animal Care and Use Committee of Xuzhou Medical University (Permit Number 2013-AN-65). The acquisition of wild *Pseudorasbora parva* in the field environment is part of a cross-sectional survey organized by Chinese Center for Disease Control and Prevention.

### Determination of *C. sinensis* infection efficiency

Q-PCR was used to determine the effectiveness of the parasite infection. The primers were designed to detect mitochondrial NADH dehydrogenase subunit 2 (*nad2*) genome of *C. sinensis* as described previously ^28^. *Nad2* primer sequences were as the following: nad2-F, GTC TGT TGA GCT TTC TCC T; nad2-R, TAA AGA CCC TGG AAA CGA GAT. For the detection of *C. sinensis* eggs, the whole liver was isolated and homogenated. Subsequently, 70% percoll density gradient was added into the sample carefully and then centrifuged at 2500 rpm for 25 min at room temperature. The cell pellet in the lowest layer was used to examine the eggs microscopically.

### Estimation of the butyrate-producing ability of the microbiota

We used a degenerate real-time PCR approach to quantify phylogenetically different butyrate-producing bacteria based on the detection of butyryl-coenzyme A (CoA) CoA transferase genes as described previously ^29^. The primer sequences were as the following: BCoATscrF (GCIGAICATTTCACITGGAAYWSITGGCAYATG) and BCoATscrR (CCTGCCTTTGCAATRTCIACRAANGC).

### Histological assays

5 weeks after parasite infection, the mice were sacrificed and the livers and extrahepatic bile ducts were removed and fixed overnight with 4% paraformaldehyde. After dehydration by gradient ethanol treatment, samples were embedded in the paraffin. After slicing, the slices were stained with haematoxylin and eosin, Alcian Blue or Masson’s trichrome, respectively. The primary antibodies Ki67 were purchased from Abcam (Cambridge, UK).

### Determination of hydroxyproline content

The concentration of hepatic hydroxyproline was determined as described in our recent publication ^30^. Briefly, frozen liver tissue was heated in 6 mM HCl at 120 °C for 4 h. After centrifugation, the supernatant was mixed with chloramine-T solution (1.4% chloramine-T, 10% N-propanol, and 80% citrate-acetate buffer) for 20 min at room temperature and followed by adding Ehrlich’s solution at 65 °C for 20 min. The absorbance at 550 nm was then measured, and the concentration was determined by the standard curve created by cis-4-hydroxy-L-proline.

### Fecal DNA Extraction

Cecal contents were collected, snap-frozen and stored at −80 °C. DNA isolation, 16S rRNA gene sequencing and bioinformatic analysis was done by Vazyme Biotech Co., Ltd., Nanjing, China. Briefly, DNA was isolated using E.Z.N.A. Stool DNA Kit (Omega Bio-Tek, Norcross, GA, USA) according to the manufacturer’s instructions. Fecal DNA samples were amplified by PCR using barcoded primer pairs targeting the V3-V4 region of 16S rRNA gene using 341F/R806 primer sets. Gel-purified PCR amplicons were sequenced using Illumina MiSeq sequencing system. The resulting bacterial sequence fragments were first clustered into Operational Taxonomic Units (OTUs) and aligned to microbial genes with 97% sequence similarity from RDP and NT-16S databases. Bacterial taxa summarization and rarefaction analyses of microbial diversity or compositional differences (dissimilarity value indicated by Unweighted UniFrac Distance) were then calculated and PCoA plots indicating compositional difference were generated accordingly with the Vegan package in R software. The data has been deposited in NCBI (accession number SRP132115).

### RNA sequencing and differential expression analyses

RNA from liver tissues were first isolated by disrupting the tissues in TRIzol (Invitrogen, Grand Island, NY, USA) using a tissue lyser and then further purified using the RNeasy mini kit (Qiagen, Germantown, MD, USA) including the on-column DNase digestion. Total RNA concentration and RNA integrity of samples was determined using the NanoDrop (Thermo Scientific, Grand Island, NY, USA) and Agilent 2100 Bioanalyzer. Further downstream library constructions were performed by Vazyme Biotech Co., Ltd., Nanjing, China. The libraries were multiplexed and sequenced on Illumia HiSeq sequencer. Transcriptional level expression analyses were performed using streamlined process with HISAT2 and cufflinks-2.2.1. The data has been deposited in NCBI (accession number GSE112003).

### Organoids experiment

The biliary organoids culture was established following a previously established protocol ^27^. Briefly, EHBD tissues from WT mice were cut into small pieces and incubated with dissociation buffer containing RPMI with 4% FCS, 1mg/ml collagenase D, 0.5mg/ml dispase, and 40 µg/ml DNase at 37°C for 30 min. Isolated cells were mixed with Matrigel and seeded onto plates overlaid with culture medium containing Ad-DMEM/F12 supplemented with 1X B27and 1X N2, 1 mM N-acetylcysteine, 10 mM nicotinamide, 50 ng/ml EGF, 1μg/ml Rspo1, 100 ng/ml Noggin, 100 ng/ml FGF10, and 500ng/ml Wnt3a,100 U/mL penicillin, 100 μg/ml streptomycin. To test the effect of butyrate, 7-9 days after initial biliary organoids culture, the organoids were further treated with 1µg/ml rmIL33 and 10mM sodium butyrate for another two days. For Ki67 immunostaining, the biliary organoids were fixed in 4% paraformaldehyde at RT for 30min, then treated with 0.1% Triton X-100 at RT for 30min, followed by blocking with 5% BSA at RT for 1h, first antibody (Ki67, Abcam, Cambridge, UK) incubation at 4°C overnight, secondary antibody (IgG H&L Alexa Fluor® 488, ThemoFisher, Waltham, MA USA) at RT for 1.5h and DAPI 5min.

### 3,5-diethoxycarbonyl-1,4-dihydrocollidine (DDC)-induced cholangiopathy model

The model was established following a previously published protocol ^3^. The animal diet containing 0.1% DDC (Sigma-Aldrich China, Shanghai, PR China) was prepared by Beijing Keao Xieli Feed Co., Ltd., Beijing. 6-8-week-old mice were fed with 0.1% DDC-containing food for total of 4 weeks. For butyrate treatment, 10g/l sodium butyrate was added in the drinking water and refreshed every other day during DCC challenge.

### Statistical Analysis

The data are shown as mean values ± standard error of the mean (SEM). Differences between multiple groups were compared using one-way ANOVA with post-hoc Bonferroni analysis. A Student’s *t*-test was used for comparisons between two groups. A *P*-value < 0.05 was considered significant.

## Author Contributions

Yugang Wang, Kuiyang Zheng, and Renxian Tang conceived and designed the experiments. Wenli Liu, Chao Yan, Bo Zhang, Renjin Chen, and Yugang Wang conducted the experiments and analyzed the data. Qian Yu, Xiangyang Li, Yuzhao Zhang, Hui Hua, Yanxia Wei, Yanbo Kou, Zhuanzhuan Liu helped for collection of *C. sinensis*, oral gavage and tissue harvest and processing. Yugang Wang wrote the paper. All authors discussed the results and commented on the manuscript.

## ACKNOWLEDGEMENTS

Project support was provided in part by the National Natural Science Foundation of China (Grant Nos. 81770853, 81702027), the Priority Academic Program Development of Jiangsu Higher Education Institutions (PAPD) in the year of 2014 (Grant No. KYLX14-1448), Natural Science Foundation of the Higher Education Institutions of Jiangsu Province, China (Grant Nos 16KJB310016), the Starting Foundation for Talents of Xuzhou Medical University (Grant No. D2016029). YGW is a specially appointed professor by Universities in Jiangsu Province.

## Data Availability

The datasets generated during the current study are available in the NCBI repository, [https://www.ncbi.nlm.nih.gov/geo/query/acc.cgi?acc=GSE112003; https://www.ncbi.nlm.nih.gov/sra/SRP132115].

## Supplementary Material

**Supplementary Figure S1.**
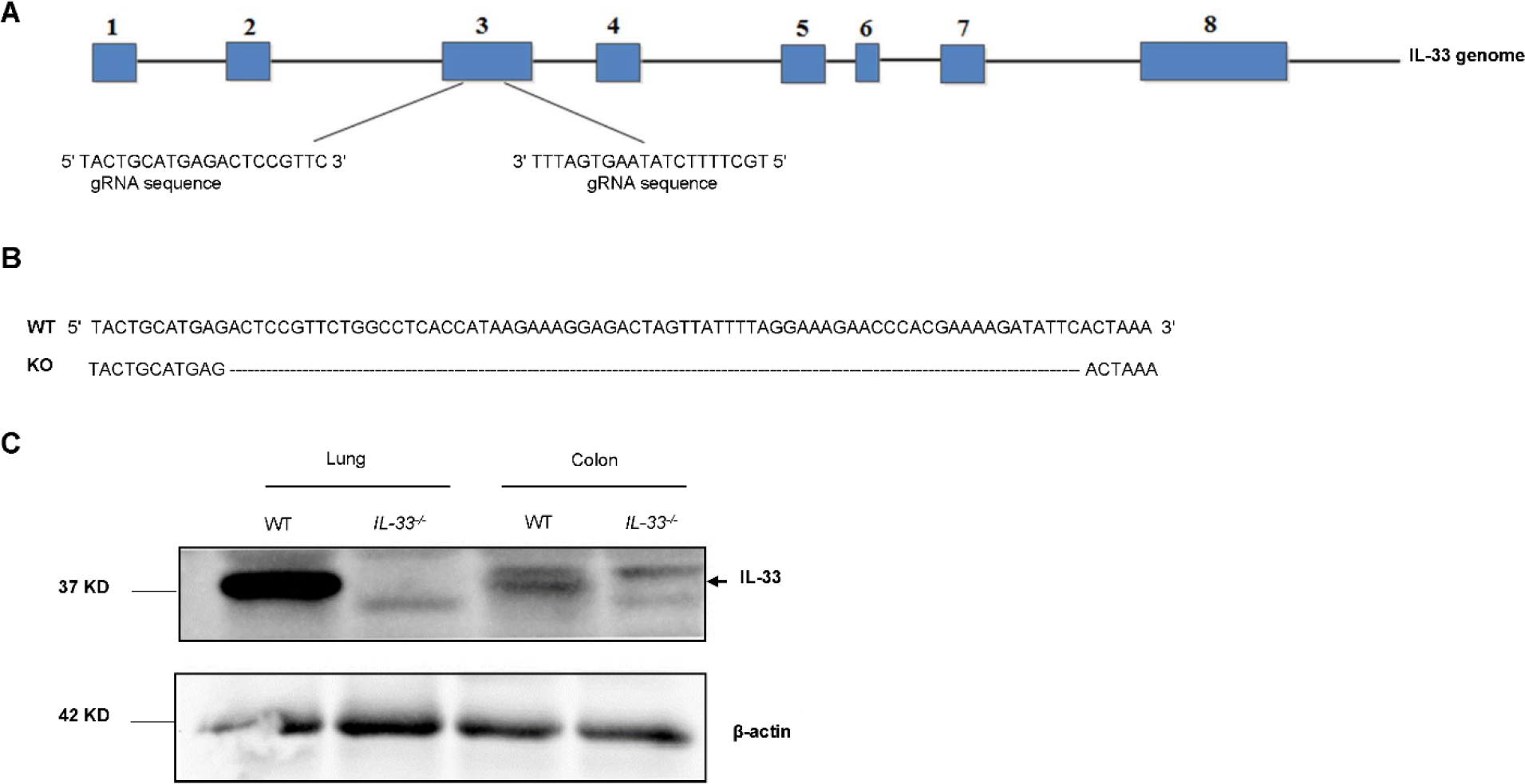
Generation of *IL-33^-/-^* mice by CRISPR/Cas9-mediated genome editing. **(A)** The targeted IL-33 locus region. The two gRNAs matching the sequences flanking the exon 3 were shown. **(B)** The mutated sequence of the knockout allele was determined by DNA sequencing and aligned with the WT allele. **(C)** IL-33 protein expression analysis by Western Blot of the colon and lung tissues from the WT and *IL-33^-/-^* mice as indicated. The blots are from different gels loaded with the same samples, and only part of the blots that close the predicted molecular sizes of the relevant proteins were cropped out for further downstream immunoblotting processing.

**Supplementary Figure S2.**
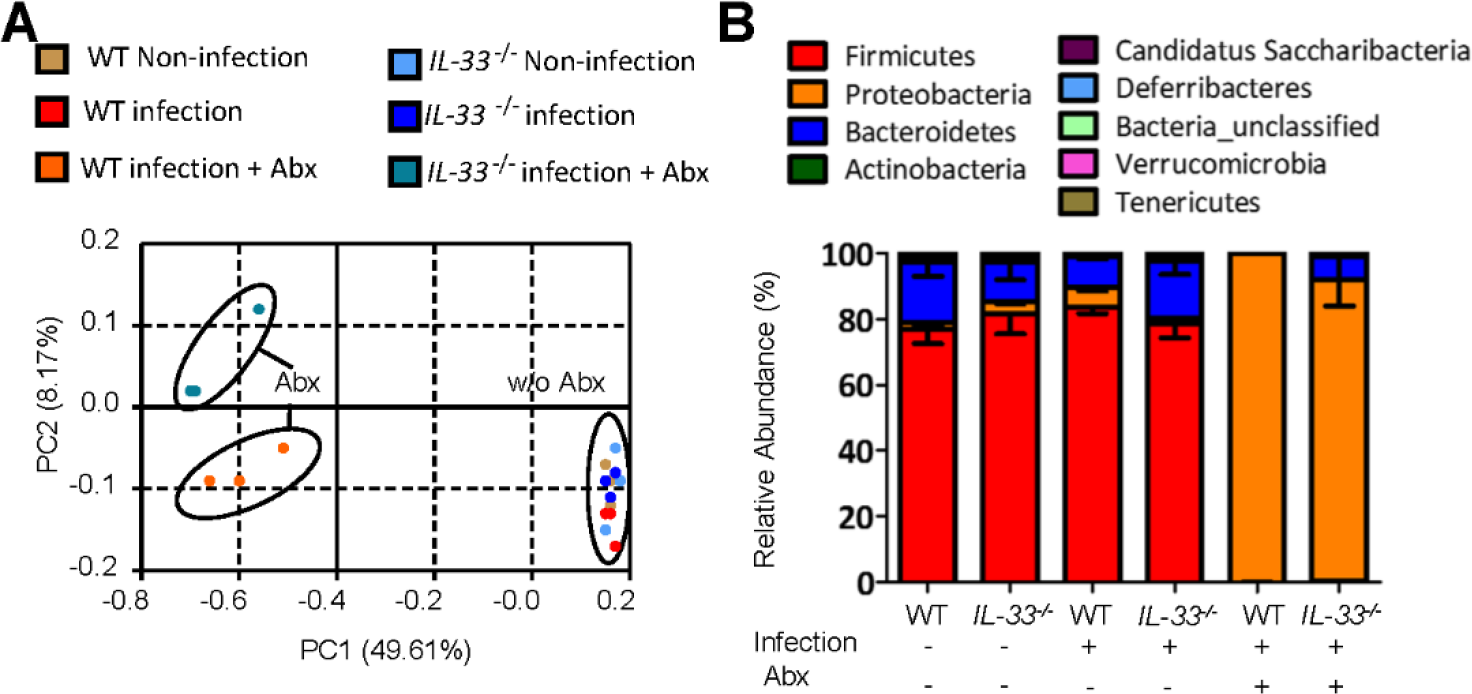
Cecal bacterial 16S rRNA sequencing analysis after abx treatment. **(A)** Unweighted UniFrac principal coordinate analysis (PCoA) showing microbiota compositional differences between antibiotics-treated and untreated mice. Each symbol represents one mouse. **(B)** Microbiota compositions at the phyla level of the indicated mice (n=3 for each, the values were expressed as mean ± SEM).

**Supplementary Figure S3.**
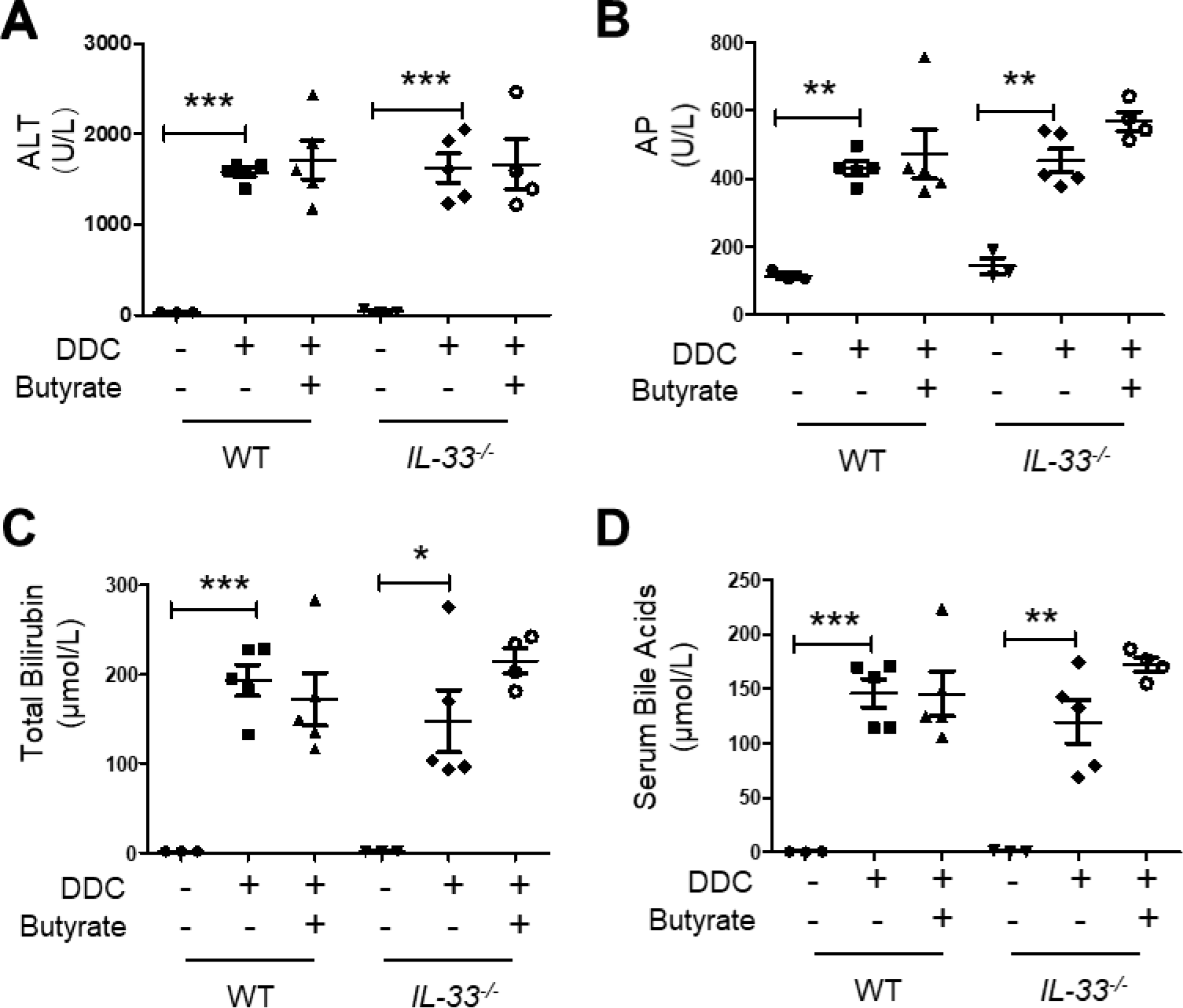
Butyrate cannot prevent DDC-induced liver injury. **(A-D)** Serum biochemistry of 4-week DDC-intoxicated mice. (A) ALT concentration; (B) Alkaline phosphatase concentration; (C) Total bilirubin concentration; (D) Serum bile acids concentration.

